# Subtelomere-specific condensed chromatin is regulated by three different histone modifications

**DOI:** 10.1101/2024.10.03.616439

**Authors:** Miho Osaki, Atika Nurani, Nanoka Asano, Yoko Otsubo, Junko Kanoh

## Abstract

In fission yeast, telomere-adjacent subtelomeres form a subtelomere-specific condensed chromatin structure, referred to as a knob, requiring histone H2A-S121 phosphorylation-dependent localization of Sgo2 at subtelomeres during interphase. However, the mechanism underlying specific Sgo2 localization in subtelomeres remains unclear. Our genetic screen identified Nts1, a histone deacetylase complex component, as a regulator of Sgo2 localization. Nts1 localized to subtelomeres during interphase and influenced histone H4 acetylation. The deletion of both Nts1 and Set2, a histone H3-K36 methyltransferase, led to the loss of Sgo2 at subtelomeres. These findings indicate that H4 deacetylation and H3-K36 methylation redundantly determine Sgo2 localization under H2A-S121 phosphorylation.

## Introduction

Telomeres are specialized chromatin structures with species-specific tandem repeat DNA at the ends of eukaryotic linear chromosomes. They contribute to various biological activities, such as protection of chromosome ends, the regulation of cell senescence, and chromosome dynamics during mitosis and meiosis (Chikashige et al. 1994; de Lange 2009; Fujita et al. 2012). Subtelomeres, telomere-adjacent domains, possess DNA sequences that are distinct from those of telomeres, but show conservation within species, with mosaics of common segments (Louis 1995; Linardopoulou et al. 2005; Oizumi et al. 2021).

In the fission yeast *Schizosaccharomyces pombe* (*S. pombe*), subtelomeres are composed of two distinct regions. The telomere-adjacent region, approximately 50 kb in length, contains subtelomeric homologous (SH) sequences with various common segments. In contrast, the telomere-distal region, also about 50 kb long, referred to as the subtelomeric unique (SU) sequence, rarely contains common sequences between subtelomeres (Tashiro et al. 2017; Oizumi et al. 2021). Furthermore, *S. pombe* subtelomeres form two distinct condensed chromatin structures. The SH regions preferentially form heterochromatin containing methylated histone H3-K9 (H3-K9me), established by telomere- and subtelomere-associated proteins and the RNA interference machinery, which recognize telomere repeats and a part of *tlh1-4* genes within the SH sequences, respectively (Cam et al. 2005; Kanoh et al. 2005; Sugiyama et al. 2007). In contrast, the SU regions, lacking significant sequence conservation across subtelomeres, typically form a condensed chromatin structure, called a knob, specifically during interphase (Matsuda et al. 2015) (Fig. 1A).

**Figure 1.**
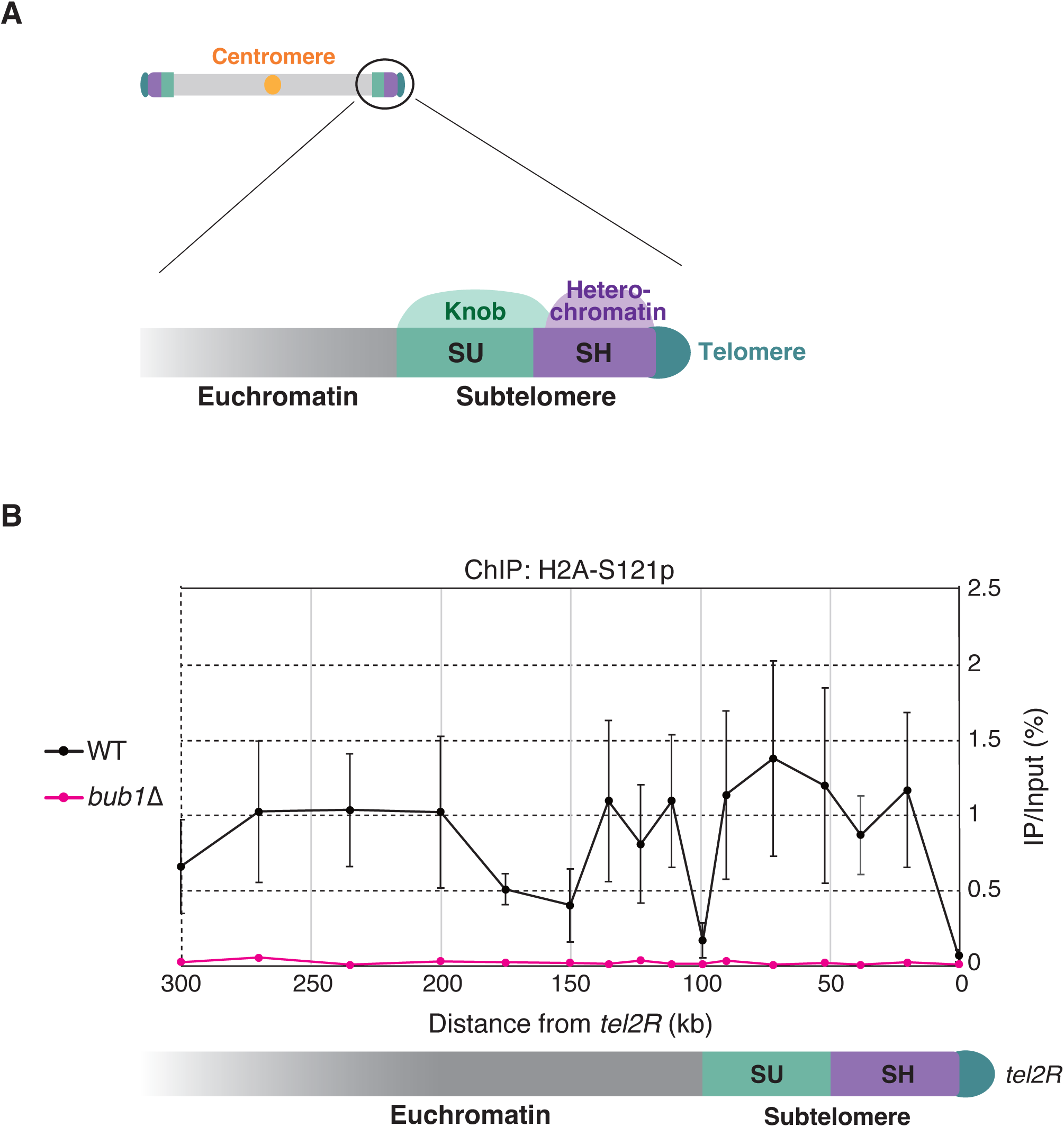
Chromatin structures of *S. pombe* subtelomeres. *(A)* Schematic illustration of *S. pombe* subtelomeres that form two distinct condensed chromatin structures. A semicircle in dark green and boxes in purple and light green represent telomeres, SH and SU regions in subtelomeres, respectively. H3K9me-dependent heterochromatin is formed within the SH region, whereas Sgo2-dependent knob is formed preferentially in the SU region. *(B)* The phosphorylation of H2A-S121 is not limited to the subtelomeres. ChIP analyses of H2A-S121p levels in the wild-type (black) and *bub1*Δ strains (magenta). X-axis shows the distance from the telomere of the right arm of chromosome 2 (*tel2R*). The recovery of immunoprecipitated DNA relative to total input DNA was measured by quantitative PCR. Error bars indicate the s.d. (*N* = 3 biologically independent experiments). Note that the values at the SH region represent the average values for all SH regions in chromosomes 1 and 2.

The Shugoshin 2 (Sgo2) protein, which is localized at pericentromeres during M phase and contributes to precise chromosome segregation, is recruited to both the SH and SU regions during interphase and plays a crucial role in knob formation (Kawashima et al. 2007; Kawashima et al. 2010; Tashiro et al. 2016). The deletion of Sgo2 causes the de-repression of subtelomeric genes and abnormal DNA replication timing of subtelomeres in S phase; thus, the knob structure and/or Sgo2 itself is important for these regulatory effects (Tashiro et al. 2016).

Little is known about how Sgo2 is recruited to the subtelomeres in interphase. The chromatin association of Sgo2 depends mostly on the Bub1-mediated phosphorylation of histone H2A-S121; thus, Bub1 deletion results in the dissociation of Sgo2 from the subtelomeres and pericentromeres (Kawashima et al. 2010; Tashiro et al. 2016). Set2, a methyltransferase for histone H3-K36, also regulates the subtelomeric localization of Sgo2 (Kizer et al. 2005; Morris et al. 2005). Set2 deletion causes the dissociation of the greater part of Sgo2 from the subtelomeres (Tashiro et al. 2016). However, detailed molecular mechanisms underlying Sgo2 recruitment to subtelomeres and the specific Sgo2 localization at subtelomeres during interphase remain unclear.

In this study, we performed a genetic screen for mutants that show the de-repression of subtelomeric genes, as observed in the *sgo2* mutant, to identify regulators of Sgo2 recruitment to subtelomeres during interphase. Some of the mutants contained mutations in the *nts1* gene, which encodes a subunit of a histone deacetylase (HDAC) (Zilio et al. 2014). We show that phosphorylation of H2A-S121 is not sufficient for Sgo2 recruitment to subtelomeres and that Nts1 and Set2 redundantly regulate the subtelomeric localization of Sgo2. Thus, three distinct histone modifications, phosphorylation, methylation, and deacetylation, cooperatively promote the Sgo2 association with subtelomeres (i.e., knob formation) during interphase.

## Results and Discussion

### Phosphorylation of histone H2A-S121 and methylation of histone H3-K36 are not confined to subtelomeres during interphase

The phosphorylation of histone H2A-S121 (H2A-S121p) by Bub1 is crucial for the association of Sgo2 with subtelomeres and centromeres in vegetatively growing cells (Kawashima et al. 2010; Tashiro et al. 2016). To investigate the regulatory mechanisms for the subtelomere-specific localization of Sgo2 in interphase, we first determined whether H2A-S121p is confined to subtelomeres in this phase. Using chromatin immunoprecipitation (ChIP) assays, we detected H2A-S121p not only at the subtelomere but also at the outer euchromatin region (Fig. 1B).

The methylation of histone H3-K36 (H3-K36me) is coupled with transcription (Kizer et al. 2005; Morris et al. 2005); thus, this modification is detected widely in the chromosomes (Dutrow et al. 2008). However, H3-K36me levels are repressed in SU regions by the subtelomeric localization of Sgo2 (Dutrow et al. 2008; Tashiro et al. 2016). These findings indicate that H2A-S121p and H3-K36me alone do not restrict Sgo2 localization to subtelomeres during interphase (i.e., additional factors are required).

### Screen for mutants that are defective in marker gene repression in SU regions

To identify novel factors that contribute to the subtelomeric localization of Sgo2, we used a genetic screening approach. When Sgo2 is not localized to subtelomeres, gene expression at SU regions in subtelomeres becomes derepressed probably due to the lack of knob structure formation (Tashiro et al. 2016). Accordingly, we screened for mutants with derepressed gene expression in SU regions, resembling the phenotype in the *sgo2*Δ strain. We first inserted three marker genes, *ade6*^+^, *his5*^+^, and *ura4*^+^, into the SU regions of three different subtelomeres (*subtel1R*, *subtel2L*, and *subtel2R*) respectively to prevent the identification of mutations in the metabolic pathway involving a single marker gene, when only one is used (Fig. 2A). Next, the parent strains were treated with nitrosoguanidine for random mutagenesis. We screened approximately 3,156,000 strains and obtained 483 mutants that grew well on the marker gene-selective medium (SD-AHU: SD w/o adenine, histidine, and uracil, but w/ leucine and lysine), as the *sgo2*Δ strain did (Fig. 2B). We selected 30 strains that grew particularly well on SD-AHU medium and analyzed the DNA sequence of the *sgo2*^+^ gene region. Mutations in *sgo2*^+^ were detected in 11 strains, indicating that this screening worked properly. Of note, we did not detect *bub1* mutations.

**Figure 2.**
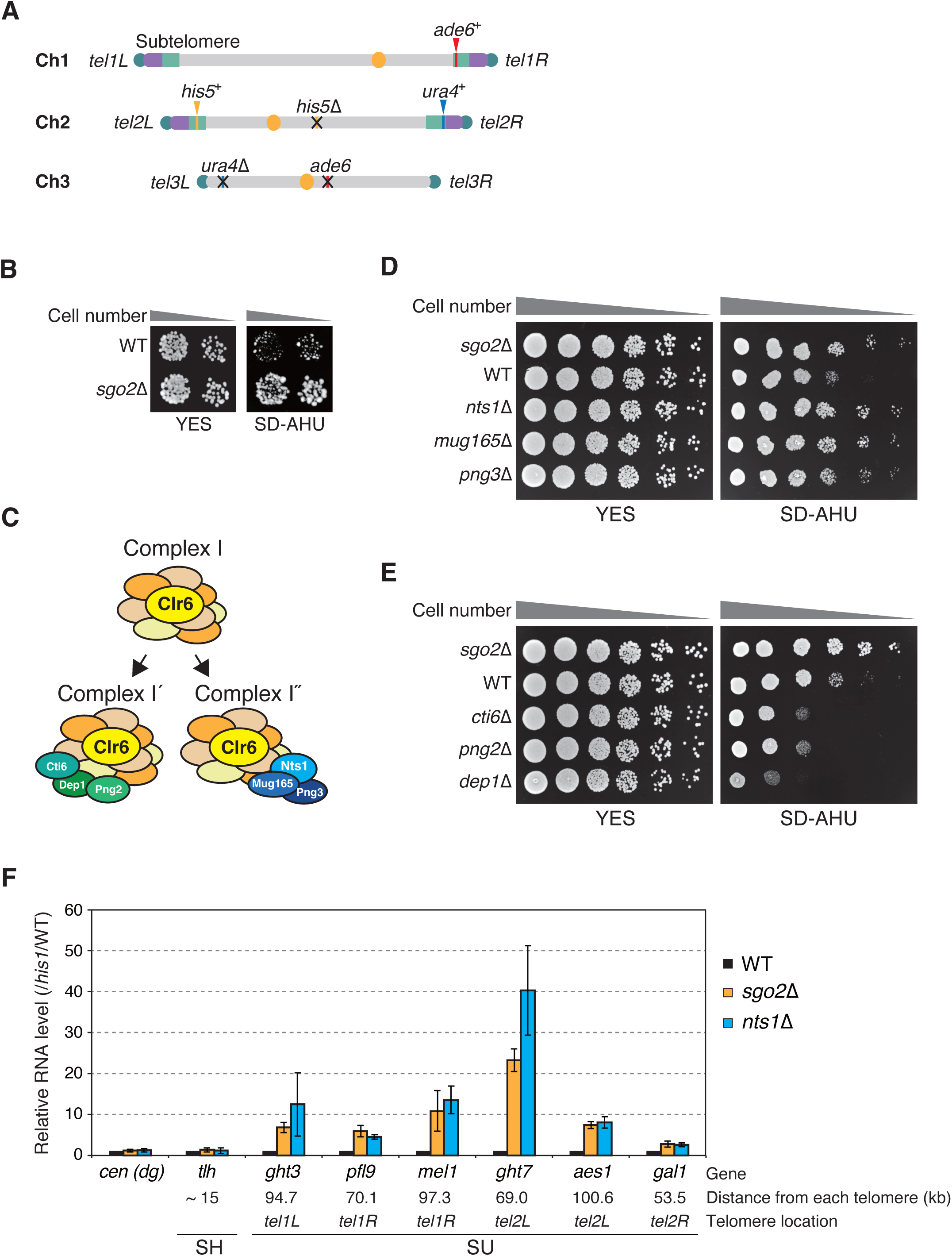
Repression of genes in SU regions requires complex I” of Clr6-HDAC. *(A)* Genetic backgrounds of the parental strain (MO4434) used for screening. Marker genes *ade6*^+^, *his7*^+^, and *ura4*^+^ were inserted into the SU regions of *subtel1R*, *subtel2L*, and *subtel2R*, respectively. Endogenous versions of these genes were deleted from the strain. Note that it is currently unknown whether the strain MO4434 has subtelomeric sequences in chromosome 3. *(B)* Better growth of *sgo2*Δ cells than wild-type cells on the selective medium (SD-AHU) for the marker genes inserted at SU regions. Cells were spotted on YES (non-selective complete medium) and SD-AHU (lacking adenine, histidine, and uracil) plates at 32°C for 2 days. *(C)* Schematic illustration of complexes I, Í, and I” with Clr6. Cti6, Png2, and Dep1 are components of complex Í, and Nts1, Mug165, and Png3 are components of I”. *(D)* Cell growth assays of the wild-type, *sgo2*Δ, *nts1*Δ, *mug165*Δ, and *png3*Δ strains. Dilution series of cells were spotted on YES and SD-AHU plates at 32°C for 2 days. *(E)* Cell growth assays of the wild-type, *sgo2*Δ, *cti6*Δ, *png2*Δ, and *dep1*Δ strains. Dilution series of cells were spotted on YES and SD-AHU plates at 32°C for 2 days. *(F)* Nts1 is required for subtelomeric gene repression. Transcript levels of subtelomeric genes and centromeric repeats (*dg*) in wild-type, *sgo2*Δ, and *nts1*Δ cells were analyzed by quantitative RT-PCR. Locations of genes are shown below. Each value was normalized first to that of *his1*^+^, and then to the wild-type value. Error bars indicate the s.d. (*N* = 3 biologically independent experiments).

### Complex I” of Clr6-HDAC is required for gene repression in SU regions

Next, we analyzed the mutant strains in which no mutations were detected in *sgo2*^+^, indicating that mutations other than those in *sgo2*^+^ and *bub1*^+^ caused derepression of the marker genes inserted in SU regions. After producing a diploid strain with the mating-type genotype *h*^-^/*h*^-^ by cell fusion of each mutant with strains where the *natMX6* marker gene (Hentges et al. 2005) was inserted into one of the three chromosomes, we artificially induced haploidization using thiabendazole (TBZ) (i.e., without meiotic recombination) to identify which chromosome harbored the critical mutation for marker gene derepression. We then inserted the *kanMX6* marker gene (Bahler et al. 1998) at various loci on the identified chromosomes and determined the chromosomal region containing the critical mutation by serial tetrad analyses. Five mutant strains had mutations in *nts1*^+^ and two had mutations in *mug165*^+^. Both Nts1 and Mug165 are components of complex I”, which interacts with one of the core complexes with Clr6 (a class I HDAC), complex I; complex I” also contains its specific component, Png3 (Grewal et al. 1998; Bjerling et al. 2002; Nicolas et al. 2007; Shevchenko et al. 2008; Zilio et al. 2014) (Fig. 2C). When we deleted the three genes encoding Nts1, Mug165, and Png3 by replacing them with marker genes respectively, all the knockout strains, like the *sgo2*Δ strain, showed better growth in the SD-AHU medium than that of the wild-type strain (Fig. 2D). In contrast, the deletion of genes encoding components of the complex Í, Cti6, Png2, and Dep1 (Nicolas et al. 2007; Shevchenko et al. 2008), had the opposite effects on cell growth in SD-AHU (Fig. 2E), suggesting that the complex I” (and not the complex Í) contributes to marker gene repression in SU regions. Subsequent analyses focused on Nts1 as a representative of complex I”.

We further investigated whether Nts1 influences the expression of intrinsic genes within the subtelomere, in addition to the marker genes. In the *nts1*Δ strain, much like in the *sgo2*Δ strain, there was no significant alteration in the RNA expression from the *dg* repeat in the centromeres or in the expression of the *tlh* genes in the SH regions of chromosomes 1 and 2 when compared with those in the wild-type strain. However, gene expression in all SU regions was notably derepressed, consistent with previous findings (Zilio et al. 2014) (Fig. 2F). These observations suggest that Nts1, like Sgo2, plays a significant role in gene repression within the SU regions.

### Nts1 and Set2 function redundantly in knob formation and Sgo2 localization at subtelomeres

The observation that Nts1 contributes to gene repression specifically in the SU region rather than in the SH region suggests that it is involved in knob formation in SU regions. To investigate this possibility, we analyzed knob formation rate in each gene-deleted strain. Knobs can be detected in interphase cells as strong histone H2A-GFP signals by fluorescence microscopy (Fig. 3A). As previously reported (Tashiro et al. 2016), the *sgo2*Δ strain exhibited almost no knob formation, and the strain lacking Set2, a methyltransferase for H3K36, displayed a reduced knob formation rate compared to that of the wild type. Similarly, the knob formation rate in the *nts1*Δ strain was lower than that of the wild type (Fig. 3B and Supplemental Fig. S1). Notably, cells deficient in both Nts1 and Set2 showed no knob formation, suggesting that Nts1 and Set2 function redundantly in knob formation (Fig. 3B and Supplemental Fig. S1). These phenotypes cannot be attributed to a decrease in Sgo2 protein levels (Supplemental Fig. S2).

**Figure 3.**
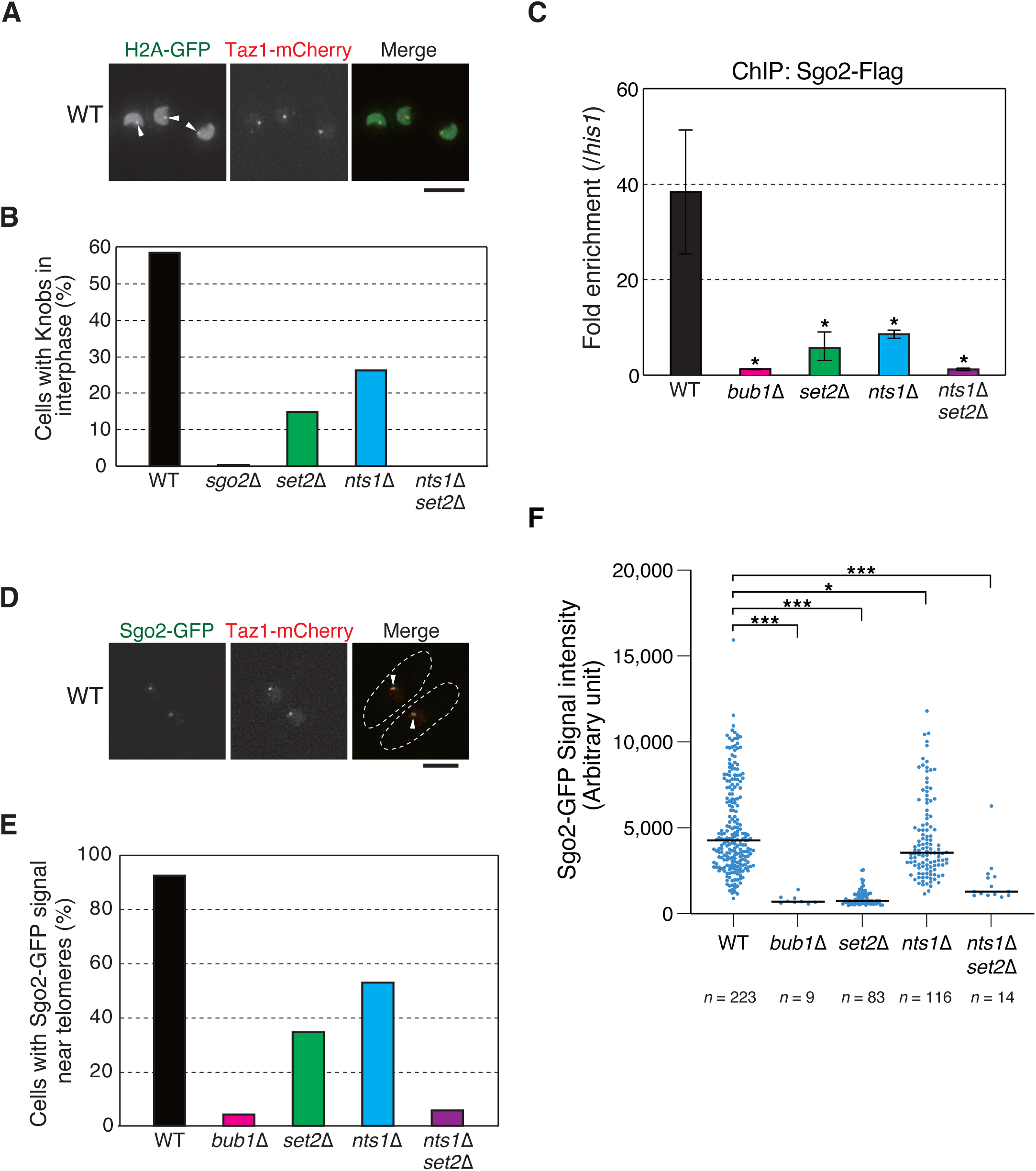
Deletion of both Nts1 and Set2 causes the dramatic loss of knob and Sgo2 at subtelomeres. *(A)* Detection of knobs as intense histone H2A-GFP signals in the wild-type strain. Arrowheads indicate knobs. Bar, 5 µm. *(B)* Frequencies of knob formation in each strain during interphase. More than 200 cells were analyzed for each strain. *(C)* ChIP analyses of Sgo2-Flag localization at *subtel2R* (51.8 kb from *tel2R*) in various deletion strains. Relative fold enrichment at *subtel2R*, normalized to the signal at the *his1*^+^ locus, is shown. Error bars indicate the s.d. (*N* = 3 biologically independent experiments). Student’s t-test (versus the wild type). 0.01 < \**p* ≤ 0.05. *(D)* Representative microscopic images of wild-type cells with Sgo2-GFP signals near telomeres (Taz1-mCherry) during interphase. Arrowheads indicate close localization of Sgo2-GFP and Taz1-mCherry. Cell outlines are indicated by broken lines. Bar, 5 µm. *(E)* Frequencies of cells with Sgo2-GFP signals near telomeres (Taz1-mCherry) during interphase. More than 200 cells were analyzed for each strain. *(F)* Intensity of each Sgo2-GFP signal near telomeres (Taz1-mCherry) during interphase. Bar, median. Mann-Whitney U test (versus the wild type). \*\*\**p* ≤ 0.001; 0.001< \*\**p* ≤ 0.01; 0.01 < \**p* ≤ 0.05.

Next, we investigated whether Nts1 is involved in the subtelomeric localization of Sgo2, which is essential for knob formation, using a ChIP assay. As reported previously, in cells deficient for Bub1, a kinase for the phosphorylation of histone H2A-S121, subtelomeric localization of Sgo2 was hardly detected at 51.8 kb from the telomere of the right arm of chromosome 2 (*tel2R*), where Sgo2 localization peaks within the subtelomere in the wild-type strain (Tashiro et al. 2016) (Fig. 3C). Consistent with the results for the knob formation rate described above, Sgo2 localization was reduced in the *set2*Δ and *nts1*Δ strains compared with the wild type but was hardly detected in the *nts1*Δ*set2*Δ strain (Fig. 3C). Similarly, in the wild-type strain during interphase, the Sgo2-GFP signal was frequently observed in proximity to the Taz1-mCherry signal (a telomeric marker), indicating its localization to the subtelomeric region. In contrast, this localization was absent in the *bub1*Δ strain (Fig. 3D and E, and Supplemental Fig. S3). In the *set2*Δ and *nts1*Δ strains, the Sgo2-GFP signal near telomeres was partially reduced compared to the wild type. However, in the *nts1*Δ*set2*Δ double mutant, like the *bub1*Δ strain, this subtelomeric localization was scarcely detected (Fig. 3E and Supplemental Fig. S3). In addition, Sgo2-GFP signal intensities near telomeres decreased substantially in the *bub1*Δ, *set2*Δ, and *nts1*Δ*set2*Δ strains but decreased slightly in the *nts1*Δ strain (Fig. 3F). These findings indicated that Nts1 and Set2 contribute redundantly to the subtelomeric localization of Sgo2.

### Nts1 accumulates at subtelomeres during interphase in a Sgo2-dependent manner

To explore the mechanism by which Nts1 acts on subtelomeres, we investigated its localization. Since ChIP assays for Nts1 were unsuccessful, we analyzed the localization of Nts1-GFP in living cells by fluorescence microscopy. In interphase cells, a distinct Nts1-GFP signal was frequently detected near telomeres (Taz1-mCherry); however, such a signal was not observed in cells during M phase (Fig. 4A). This suggests that Nts1 accumulates in the vicinity of telomeres, likely at subtelomeres, during interphase.

**Figure 4.**
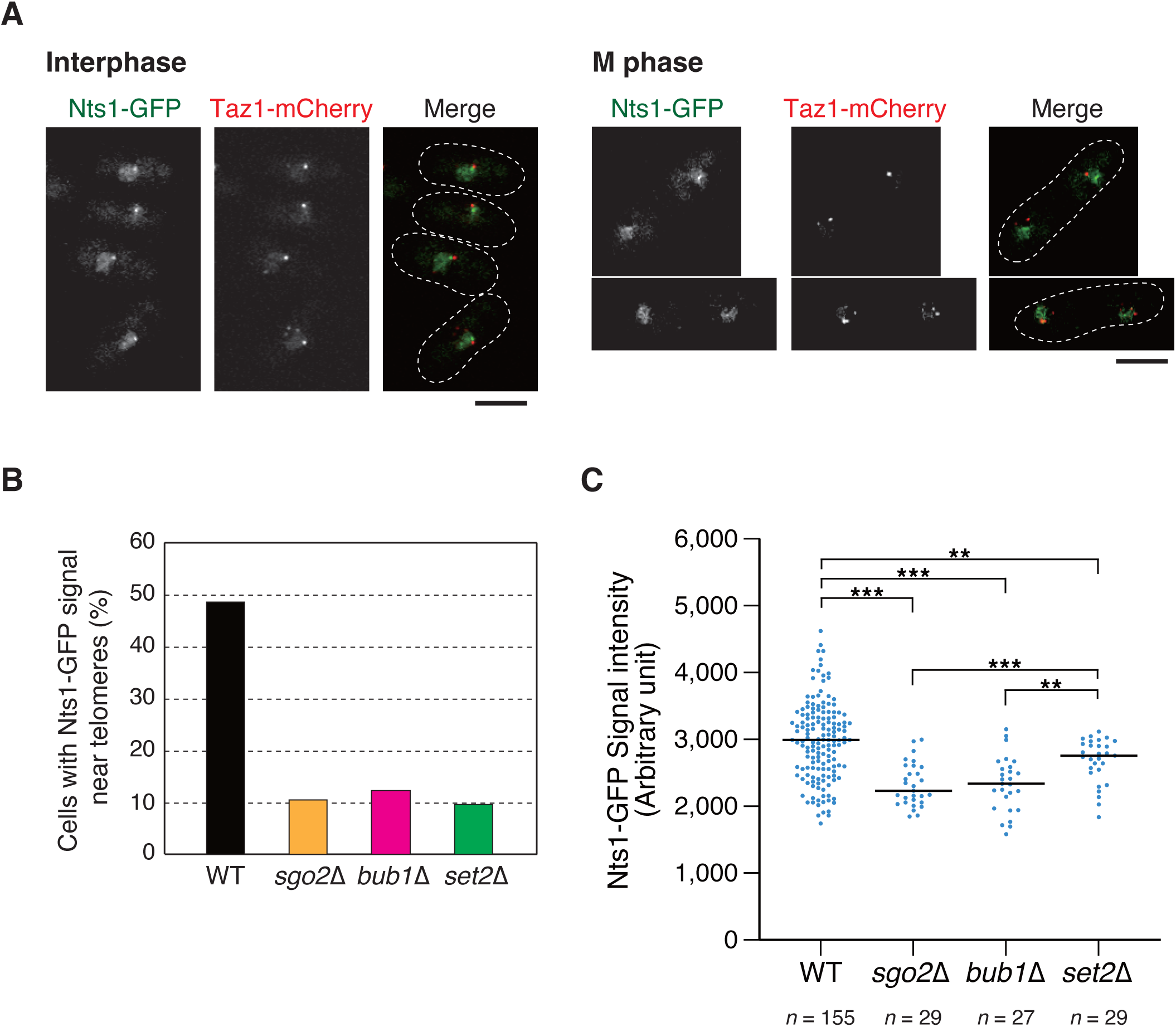
Nts1 acts on subtelomeres during interphase in a Sgo2-dependent manner. *(A)* Representative microscopic images of wild-type cells expressing Nts1-GFP and Taz1-mCherry (indicating telomeres) in interphase (left) and M phase (right). Cell outlines are indicated by broken lines. Bar, 5 µm. *(B)* Frequencies of cells with Nts1-GFP signals near telomeres (Taz1-mCherry) during interphase. More than 200 cells were analyzed for each strain. *(C)* Intensity of each Nts1-GFP signal near telomeres (Taz1-mCherry) during interphase. Bar, median. Mann-Whitney U test (versus the wild type or *set2*Δ). \*\*\**p* ≤ 0.001; 0.001 < \*\**p* ≤ 0.01.

Interestingly, the telomere-proximal (likely subtelomeric) localization of Nts1-GFP was decreased in *sgo2*Δ, *bub1*Δ, and *set2*Δ strains, although the protein expression levels of Nts1-GFP in these strains were comparable to that in the wild type (Fig. 4B and Supplemental Fig. S4). Moreover, the Nts1-GFP signal intensities in the *sgo2*Δ and *bub1*Δ strains were significantly weaker than those in the wild-type or *set2*Δ strains, whereas only mild effects were observed in the *set2*Δ strain (Fig. 4C). These results suggest that the subtelomeric localization of Sgo2 is required for the efficient accumulation of Nts1 at subtelomeres; that is, the subtelomeric localizations of Nts1 and Sgo2 are interdependent.

### Nts1 contributes to the deacetylation of histone H4 in the SU region

The Clr6-HDAC complex deacetylates histones H3 and H4, contributing to gene repression (Grewal et al. 1998; Bjerling et al. 2002; Wiren et al. 2005; Hansen et al. 2011; Wang et al. 2023). To investigate whether complex I” of Clr6-HDAC, which includes Nts1, is involved in histone deacetylation in subtelomeres, we focused on the acetylation of histone H4 at K5 and K12, which shows lower acetylation in subtelomeres than in euchromatin regions (Wiren et al. 2005; Buchanan et al. 2009). ChIP analyses revealed that the acetylation levels of H4-K5 and H4-K12 were higher in the *nts1*Δ strain than in the wild type, especially in the SU region (Fig. 5A and B). Furthermore, consistent with the co-dependence of the subtelomeric localization of Sgo2 and Nts1, similar effects on acetylation levels were observed in the *bub1*Δ strain, which does not lack Nts1 (Fig. 5A and B). These effects could be due to the insufficient association of Nts1 in the SU region in the *bub1*Δ strain. Taken together, these results indicated that complex I” of Clr6-HDAC contributes to the deacetylation of histones, such as H4 at K5 and K12, in the SU region during interphase to promote Sgo2 localization and knob formation.

**Figure 5.**
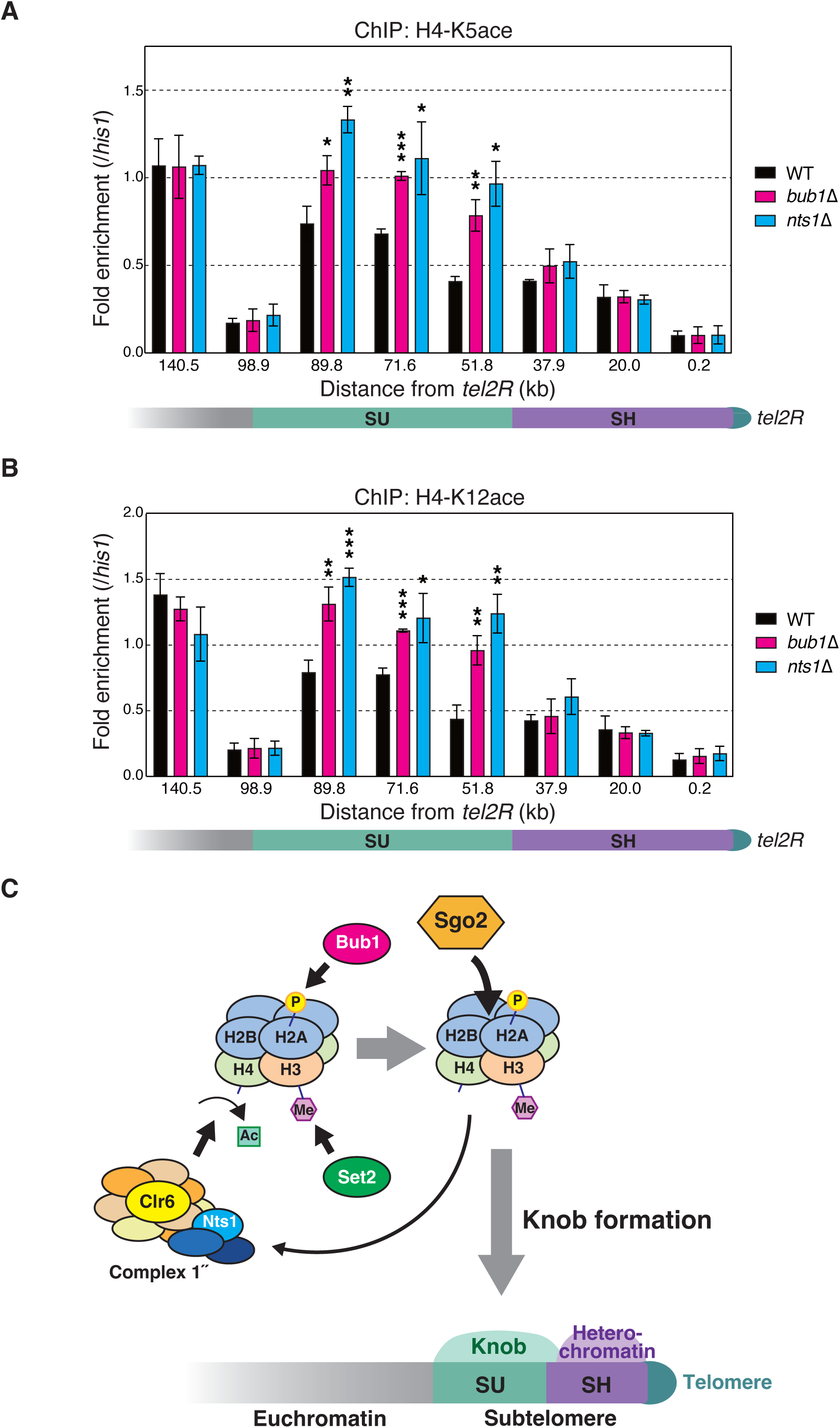
Nts1 contributes to the deacetylation of histone H4 at K5 and K12 in the SU region. *(A)* ChIP analyses of H4-K5ace levels at *subtel2R* in each strain. Relative fold enrichment at *subtel2R*, normalized to the signal at the *his1*^+^ locus, is shown. Error bars indicate the s.d. (*N* = 3 biologically independent experiments). Student’s *t*-test (versus the wild type). \*\*\**p* ≤ 0.001; 0.001< \*\**p* ≤ 0.01; 0.01 < \**p* ≤ 0.05. Note that the values at the SH region represent the average values of all SH regions. *(B)* ChIP analyses of H4-K12ace levels at *subtel2R* in each strain. Relative fold enrichment at *subtel2R*, normalized to the signal at the *his1*^+^ locus, is shown. Error bars indicate the s.d. (*N* = 3 biologically independent experiments). Student’s *t*-test (versus the wild type). \*\*\**p* ≤ 0.001; 0.001< \*\**p* ≤ 0.01; 0.01 < \**p* ≤ 0.05. Note that the values at the SH region represent average values of all SH regions. *(C)* Model of the mechanisms underlying knob formation in the SU region at subtelomeres. During interphase, complex 1” of Clr6-HDAC with Nts1 accumulates at subtelomeres and deacetylates histone H4 at K5 and K12, Set2 methylates histone H3-K36, and Bub1 phosphorylates histone H2A-S121. These changes in histone modifications promote subtelomeric localization of Sgo2 and facilitate knob formation. Sgo2 at subtelomeres further promotes the Nts1 (complex 1” of Clr6-HDAC) localization at subtelomeres via a positive feedback mechanism.

In this study, we discovered new regulatory mechanisms for the formation of the subtelomere-specific knob structure. We showed that three histone modification changes, the phosphorylation of H2A-S121 by Bub1, deacetylation of H4 via complex I” of Clr6-HDAC, and methylation of H3-K36 by Set2, promote Sgo2 localization and knob formation at subtelomeres in interphase. Our results suggest that complex I” of Clr6-HDAC acts on the SU regions within the subtelomeres during interphase; thus, it is highly possible that H4 deacetylation at SU regions is at least one of the triggers for the subtelomere-specific localization of Sgo2 in interphase. Furthermore, the subtelomeric localizations of Sgo2 and Nts1 are mutually dependent. This positive feedback regulation may be important for timely and stable knob formation during interphase (Fig. 5C).

The simultaneous deletion of Nts1 and Set2, but not each single deletion, resulted in the almost complete loss of Sgo2 and knob formation at subtelomeres, as observed in the *bub1*Δ mutant. This finding suggested that Sgo2 has at least two distinct domains that regulate its chromatin association: one affected by H2A-S121p and another by H3-K36me and the deacetylation of histones, such as H4 at K5 and K12. In fact, the shugoshin family proteins share two conserved domains, the N-terminal coiled-coil and the C-terminal basic domains; the latter of which binds directly to H2A-S121p in nucleosomes, which is essential for the chromatin localization of Sgo2 in growing cells (Kitajima et al. 2004; Liu et al. 2015). Interestingly, Sgo2 becomes associated with subtelomeres even in *bub1*Δ cells when the cells are in the stationary phase after grown in a low-glucose medium, and this Bub1-independent subtelomeric localization of Sgo2 does not require the C-terminal basic domain; however, it does require Set2 (Kobayashi and Kawashima 2019). Moreover, the N-terminal coiled-coil domain is required for dimerization of Sgo2 and its interaction with other proteins, such as Bir1, which is required for the localization of Sgo2 at pericentromeres during M phase (Xu et al. 2009; Tsukahara et al. 2010). Thus, it is possible that the coiled-coil and/or internal domains of Sgo2 mediates the interaction with nucleosomes containing H3-K36me and deacetylated H4.

The depletion of Nts1 led to an increase in the acetylation of histone H4 primarily in the SU region (Fig. 5A and B), suggesting that complex 1” of Clr6-HDAC acts at the SU region. However, it remains unclear how complex 1” is consistently targeted to the SU regions across subtelomeres, which do not share high sequence homology. The recruitment of Sgo2 to SU regions in cells lacking the adjacent SH sequences suggests that relatively short homologous sequences between SUs may serve as markers for attracting complex 1”; alternatively, once Sgo2 is recruited to the subtelomeres by the SH sequences, it may spread and maintain its subtelomeric localization by recruiting complex 1” in a manner similar to the mechanism by which HP1 attracts histone methyltransferases to maintain heterochromatin.

## Materials and Methods

### Strains and general techniques for *S. pombe*

*S. pombe* strains used in this study are listed in Supplemental Table S1. Growth media and basic genetic and biochemical techniques used in this study were described previously (Moreno et al. 1991; Bahler et al. 1998; Forsburg and Rhind 2006). Sequences of the primer sets used to generate DNA fragments to produce deletion and carboxy-terminal epitope-tagged strains are listed in Supplemental Table S2.

### ChIP analysis

ChIP was performed as described previously (Tashiro et al. 2016) with anti-H2A-S121p (Kawashima et al. 2010), anti-H3-K9me2 (MABI0307; MAB Institute), anti-H4-K5ace (ab51997; Abcam), andti-H4-K12ace (ab46983; Abcam), anti-Flag (F3165; Sigma), anti-H3-K36me3 (MABI0333; MAB Institute) antibodies. Sequences of the primer sets for subsequent quantitative PCR are listed in Supplemental Table S3.

### Isolation of mutants defective in gene silencing at SU regions

The parental strain for mutagenesis was produced as follows. The *his5*^+^ gene was deleted by replacing it with a hygromycin resistance gene cassette (*hphMX6*) (Hentges et al. 2005) in the strain JM3257 (*h*^-^ *ade6-M210 leu1-32 ura4-D18*). This was followed by the insertion of an *ade6*^+^ gene cassette at approximately 96 kb from *tel1R* in chromosome 1, a *his5*^+^ gene cassette at approximately 72 kb from *tel2L* in chromosome 2, and a *ura4*^+^ gene cassette at approximately 52 kb from *tel2R* in chromosome 2. The resultant strain (MO4434) was mutagenized with 50 µg/mL *N*-methyl-*N*’-nitro-*N*-nitrosoguanidine (Tokyo Chemical Industry), which was sufficient for approximately 70% killing (Uemura and Yanagida 1984). Surviving cells were grown in YES liquid media for 4 h, washed with H_2_O for four times, and then incubated on an SD plate without adenine, histidine, and uracil (SD-AHU) for several days at 32°C. This screen identified 483 colonies that grew vigorously on the SD-AHU plate, like the *sgo2*Δ strain (MO4326), from approximately 3,156,000 surviving strains.

### Cell fusion and haploidization

*h*^-^ strains were fused to produce *h*^-^/*h*^-^ diploids as follows. Cells were grown in a YES liquid medium, collected by centrifugation, and then suspended in buffer A (1.2 M sorbitol, 50 mM citrate phosphate [pH 5.6], 50 mM β3-mercaptoethanol). The cell suspensions of different strains were mixed and treated with Lysing Enzymes from *Trichoderma harzianum* (Sigma-Aldrich) for 1 h at 30°C with gentle shaking. The protoplasts were washed with buffer B (1.2 M sorbitol, 30 mM Tris-HCl [pH 7.5]), and PEG buffer (30% PEG 4000, 10 mM Tris-HCl [pH 7.5], 10 mM CaCl_2_) was poured onto the pellets. After incubation at 25°C for 30 min with gentle shaking, the pellet cells were resuspended in a YES liquid medium with 1.2 M sorbitol and incubated at 32°C for overnight. The cells were grown on a selective medium plate. The resultant diploid cells were treated with 30 µg/mL TBZ (Sigma-Aldrich) in a YES liquid medium and incubated overnight at 32°C. The cells were incubated on a selective medium plate supplemented with 20 mg/L of Phloxine B (Nacalai tesque) to identify haploid cells.

### RNA analyses

Total RNA was prepared from exponentially growing cells as described previously (Tashiro et al. 2016). For reverse transcription (RT)-PCR, complementary DNA was synthesized using a High-Capacity cDNA Reverse Transcription Kit (Thermo) with random primers and analyzed by quantitative PCR using the StepOne Real-Time PCR System (Thermo). Sequences of the primer sets used for quantitative PCR are listed in Supplemental Table S3.

### Microscopy

For the observation of knobs, Sgo2, or Nts1, cells expressing Hta1 (histone H2A)-GFP, Sgo2-GFP, or Nts1-GFP, respectively, and Taz1-mCherry were grown in a YES liquid medium at 32°C to the exponential phase, and images of 13 optical sections were obtained at focus intervals of 0.3 µm using a Delta Vision microscope system (Cytiva) and deconvolved by a three-dimensional deconvolution method using SOFTWORX software (Agard et al. 1989). For the analyses of signal intensities, peak levels of each Sgo2-GFP signal were analyzed using SOFTWORX software.

## Acknowledgements

We thank S. Tashiro and S. Hatano for valuable advice on the ChIP analysis, S. Kawashima for providing anti-H2A-S121p, and all the lab members for discussion and support. This work was supported by Japan Society for the promotion of Science (JSPS) KAKENHI [JP20H03185, JP20H05388, JP21H00244, JP22H04685, JP23H02408], Ohsumi Frontier Science Foundation, and Bioscience Research Grant of Takeda Science Foundation.

## Author contributions

J.K. conceived the project and designed experiments. M.O., A.N., N.A., Y.O., and J.K. performed experiments and analyzed data. J.K. and Y.O. wrote the manuscript.

